# An rRNA fragment in extracellular vesicles secreted by human airway epithelial cells increases the fluoroquinolone sensitivity of *P. aeruginosa*

**DOI:** 10.1101/2022.04.19.488721

**Authors:** Katja Koeppen, Thomas H. Hampton, Scott A. Gerber, Young Ah Goo, Byoung-Kyu Cho, Danielle M. Vermilyea, Deborah A. Hogan, Bruce A. Stanton

## Abstract

Lung infection by antibiotic resistant strains of *Pseudomonas aeruginosa* is a well-known concern for immunocompromised hosts including people with lung diseases such as cystic fibrosis. We have previously demonstrated that extracellular vesicles (EVs) secreted by primary human airway epithelial cells (AEC) deliver miRNA let-7b-5p to *P. aeruginosa* where it suppresses biofilm formation and increases sensitivity to beta-lactam antibiotics. In this study we used RNA-seq to characterize the small RNA (sRNA) content of EVs secreted by AEC and demonstrate transfer of multiple distinct RNA fragments from EVs to *P. aeruginosa*. Bioinformatic predictions reveal that several sRNAs may target all three subunits of the fluoroquinolone efflux pump MexHI-OpmD, an effect predicted to increase antibiotic sensitivity to fluoroquinolone antibiotics. Exposure of *P. aeruginosa* to EVs resulted in a significant reduction in the protein levels of MexH (−48%), MexI (−50%) and OpmD (−35%). Moreover, EVs reduced planktonic growth of *P. aeruginosa* in the presence of the fluoroquinolone antibiotic ciprofloxacin by 20%. A *mexGHI-opmD* deletion mutant of *P. aeruginosa* phenocopied this increased sensitivity to ciprofloxacin. Finally, we found that a fragment of an 18S rRNA external transcribed spacer that was transferred to *P. aeruginosa* by EVs was sufficient to reduce planktonic growth of *P. aeruginosa* in the presence of ciprofloxacin, to reduce the minimum inhibitory concentration (MIC) of *P. aeruginosa* for ciprofloxacin by over 50%, and to significantly reduce protein levels of MexH and OpmD. In conclusion, an rRNA fragment secreted by AEC in EVs increases the ciprofloxacin sensitivity of *P. aeruginosa* by targeting and down-regulating the fluoroquinolone efflux pump MexHI-OpmD. A combination of rRNA fragments and ciprofloxacin packaged in nanoparticles or EVs may benefit patients with antibiotic-resistant *P. aeruginosa* infections.

**Author Summary:** According to the World Health Organization and the U.S. Centers for Disease Control and Prevention the development of antibiotic resistant strains of bacteria, including *Pseudomonas aeruginosa*, are a significant global threat to human health. Thus, development of new approaches to eliminate antibiotic resistant infections is required. In this study, we report that lung epithelial cells secrete extracellular vesicles (EVs) that fuse with and deliver small rRNAs to *P. aeruginosa*, and that the rRNAs increase the sensitivity of *P. aeruginosa* to the antibiotic ciprofloxacin by reducing protein levels of the drug efflux pump MexHI-OpmD. We identified one rRNA fragment that by itself significantly reduced the protein levels of MexH and OpmD and increased the ability of ciprofloxacin to kill *P. aeruginosa*. We propose that developing synthetic vesicles containing a combination of the rRNA that inhibits antibiotic efflux pumps and ciprofloxacin would benefit patients with antibiotic resistant *P. aeruginosa* infections.

## Introduction

*Pseudomonas aeruginosa* is an opportunistic bacterial pathogen that can cause chronic infections in immunocompromised hosts, leading to increased morbidity and mortality in individuals with cystic fibrosis (CF), chronic obstructive pulmonary disease (COPD), ventilator-associated pneumonia and burn wounds (1). *P. aeruginosa* is also emerging as a pathogen that commonly co-infects and exacerbates COVID-19 infections (2–5). The rise of difficult-to-eradicate, antibiotic resistant strains of *P. aeruginosa* is posing treatment challenges and increasingly becoming a concern in healthcare settings (6–9). Multidrug-resistant strains of *P. aeruginosa* are estimated to have caused about 32,600 hospital-acquired infections and 2700 deaths in the US in 2017 and have been rated as a serious threat by the CDC in a 2019 report (10). Thus, new approaches are needed to eradicate antibiotic resistant infections by *P. aeruginosa*.

Bi-directional host-pathogen communication through short RNA (sRNA) in extracellular vesicles (EVs) and the resulting regulation of the host immune response as well as bacterial phenotypes such as antibiotic sensitivity have become a research focus in recent years (11–19). We have previously shown that *P. aeruginosa* outer membrane vesicles deliver a transfer RNA (tRNA) fragment to human airway epithelial cells (AEC), suppressing the host innate immune response to infection, thereby facilitating chronic infections (11). Eukaryotic EVs have also been reported to contain tRNA fragments (20–22) and tRNA fragments regulate gene expression in eukaryotes and prokaryotes (23–27), and to play a role in various human diseases (28,29). Previous studies have primarily focused on the regulatory effects of tRNA fragments from a given organism on that same organism, rather than on inter-species crosstalk. Much less is known about the role of tRNA fragments and other sRNAs in inter-kingdom signaling between eukaryotic and prokaryotic organisms.

Besides tRNA fragments and messenger RNAs (mRNA), eukaryotic EVs contain a diverse assortment of non-coding RNA species, including long non-coding RNAs (lncRNA), microRNAs (miRNA), piwi-interacting RNAs (piRNA), Y RNAs, antisense RNAs, transfer RNAs (tRNA), tRNA fragments, and ribosomal RNAs (rRNA) (20,22,30–33). EVs as well as their miRNA cargo are involved in immune regulation and their secretion and RNA content have been shown to be dysregulated in patients with airway diseases (34–38). For example, a recent study found that increased levels of miR-223-3p or miR-451a and decreased levels of miR-27b-3p in sputum supernatants correlated with exacerbation in children with CF and infection with *Aspergillus, Haemophilus*, or *Pseudomonas*, respectively, suggesting the involvement of EV miRNAs in pulmonary exacerbation (39). Two recent studies provide a comprehensive characterization of the sRNA content of EVs secreted by AEC – one reporting changes in the sRNA content of EVs secreted by the human airway epithelial cell line A549 in response to viral infection (40), and another describing cigarette smoke induced changes in the RNA content of EVs secreted by primary small airway epithelial cells (41). Prior studies of EVs secreted by airway epithelial cells (AEC) have focused mainly on their miRNA cargo (19,42–44), whereas the function of other sRNA species contained in AEC EVs has not yet been reported. While there is a growing body of literature on the regulatory effects of EV sRNAs within a given organism, little is known about the ability of sRNAs secreted in eukaryotic EVs to regulate gene expression in prokaryotes. For example, eukaryotic miRNAs affect the growth of gut microbes through an unknown mechanism (45–47). Recently, we provided the first direct evidence that a miRNA, let-7b-5p, in EVs secreted by human airway epithelial cells, is transferred to *P. aeruginosa*, where it increases beta-lactam antibiotic sensitivity and decreases biofilm formation by reducing the abundance of the beta-lactamase AmpC and several proteins required for biofilm formation (19). Here, we report that human EVs also reduce the abundance of all three subunits of the fluoroquinolone efflux pump MexHI-OpmD and increase fluoroquinolone sensitivity, an effect mediated by an rRNA fragment secreted in EVs.

## Results

### EVs increase the fluoroquinolone sensitivity of *P. aeruginosa*

In a previous study we demonstrated that EVs secreted by AEC increased the sensitivity of *P. aeruginosa* to beta lactam antibiotics by delivering the miRNA let-7b to *P. aeruginosa* (19). To test the hypothesis that EVs may also alter the sensitivity to other antibiotics typically used to treat infections by *P. aeruginosa* (48), we examined the ability of EVs to alter the sensitivity of *P. aeruginosa* to the fluoroquinolone antibiotic ciprofloxacin. To that end, the planktonic growth of *P. aeruginosa* strain PA14 as well as the mucoid *P. aeruginosa* CF clinical isolate SMC1585 (49) were measured in the presence and absence of EVs and in the presence and absence of eight different concentrations of ciprofloxacin to determine the minimal inhibitory concentration (MIC). EVs alone had no significant effect on planktonic growth of PA14 in the absence of ciprofloxacin (Fig 1A) but reduced the MIC of ciprofloxacin by 25% (Fig 1B). EVs reduced planktonic growth of PA14 in the presence of 0.02 µg/ml ciprofloxacin (corresponding to about one third of the MIC) by 26%, compared to 0.02 µg/ml ciprofloxacin alone (Fig 1C). Likewise, EVs alone had no significant effect on planktonic growth of clinical isolate SMC1585 in the absence of ciprofloxacin (Fig 1D) but reduced the MIC of ciprofloxacin by 48% (Fig 1E). EVs reduced planktonic growth of SMC1585 in the presence of 0.02 µg/ml ciprofloxacin by 27%, compared to 0.02 µg/ml ciprofloxacin alone (Fig 1F). Full 20-hour growth curves for EV-exposed and vehicle exposed PA14 and SMC1585 in the presence and absence of 0.02 µg/ml ciprofloxacin are shown in S1 Fig. In the presence of 0.02 µg/ml ciprofloxacin there was a statistically significant difference in the growth curves of PA14 and SMC1585 with EVs compared to controls for the last 4 hours of growth. Thus, the combination of EVs and ciprofloxacin reduced the MIC of ciprofloxacin as well as the growth of PA14 and a clinical isolate of *Pseudomonas* (SMC1585).

**Fig 1.**
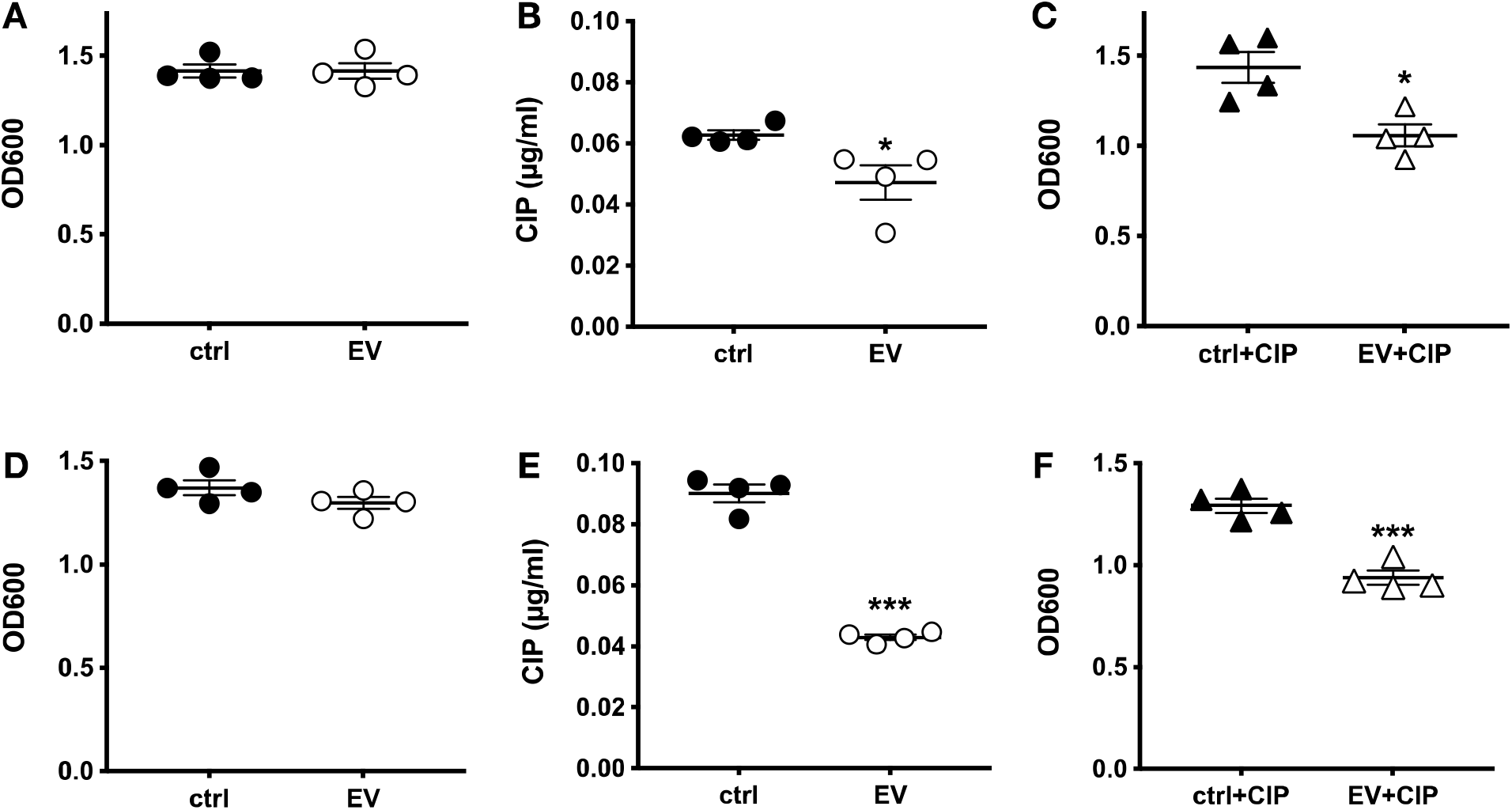
EVs increase the fluoroquinolone sensitivity of *P. aeruginosa*. EVs did not significantly affect planktonic growth (OD600) of PA14 in the absence of ciprofloxacin (CIP) after a 20-hour incubation (A), but significantly reduced the MIC of ciprofloxacin (B) as well as planktonic growth of PA14 in the presence of 0.02 µg/ml ciprofloxacin (C). EVs did not significantly affect planktonic growth (OD600) of the clinical isolate SMC1585 in the absence of ciprofloxacin after a 20-hour incubation (D), but significantly reduced the MIC of ciprofloxacin (E) as well as planktonic growth in the presence of 0.02 µg/ml ciprofloxacin (F). Statistical significance was determined using unpaired t-tests. EVs were isolated from four AEC donors and each data point is the mean of three technical replicates from each donor. ***P < 0.001, *P < 0.05.

### EVs increase the fluoroquinolone sensitivity of *P. aeruginosa* by targeting MexHI-OpmD

To begin to elucidate the mechanism whereby EVs increased the sensitivity of *P. aeruginosa* to ciprofloxacin we analyzed the proteome of *P. aeruginosa* that had been exposed to EVs or vehicle control. EVs reduced the protein levels of all three subunits of the fluoroquinolone efflux pump MexHI-OpmD (Table 1). Compared to control, EVs decreased protein levels of MexH by 48%, MexI by 50% and OpmD by 35%. Normalized peak intensities for each sample set as well as average log2 fold changes and p-values for all 1619 detected proteins in *P. aeruginosa* are provided as S1 Table. To determine whether a reduction of MexHI-OpmD alone increased fluoroquinolone sensitivity, planktonic growth experiments were performed with a *mexGHI-opmD* deletion strain of *P. aeruginosa* PAO1 and the parental PAO1 wild type strain. In the presence of ciprofloxacin EVs significantly decreased planktonic growth of wild type PAO1 (Fig 2). By contrast, in the presence of ciprofloxacin, EVs had no significant effect on the *mexGHI-opmD* deletion strain (Δ+EV). S2 Fig contains full 20-hour growth curves for the data shown in Fig 2. Taken together, the data suggest that EVs increase *P. aeruginosa* ciprofloxacin sensitivity by targeting and down-regulating the fluoroquinolone efflux pump MexHI-OpmD.

**Table 1.** EVs significantly reduce the protein levels of all three subunits of the fluoroquinolone efflux pump MexHI-OpmD.

| Uniprot ID | Locus | Name | Product | log2 FC | % decrease | P-value |
| --- | --- | --- | --- | --- | --- | --- |
| A0A0H2ZGA2 | PA14_09530 | MexH | RND efflux membrane fusion protein | -0.93 | -48 | 0.0008 |
| A0A0H2ZGB3 | PA14_09520 | MexI | RND efflux transporter | -1.01 | -50 | 0.0003 |
| A0A0H2ZF64 | PA14_09500 | OpmD | outer membrane protein | -0.62 | -35 | 0.0161 |

Protein level changes in *P. aeruginosa* exposed to EVs compared to vehicle controls. FC = fold change, N = three biological replicates of EVs secreted by different AEC donors. Statistical significance was determined with QPROT.

**Fig 2.**
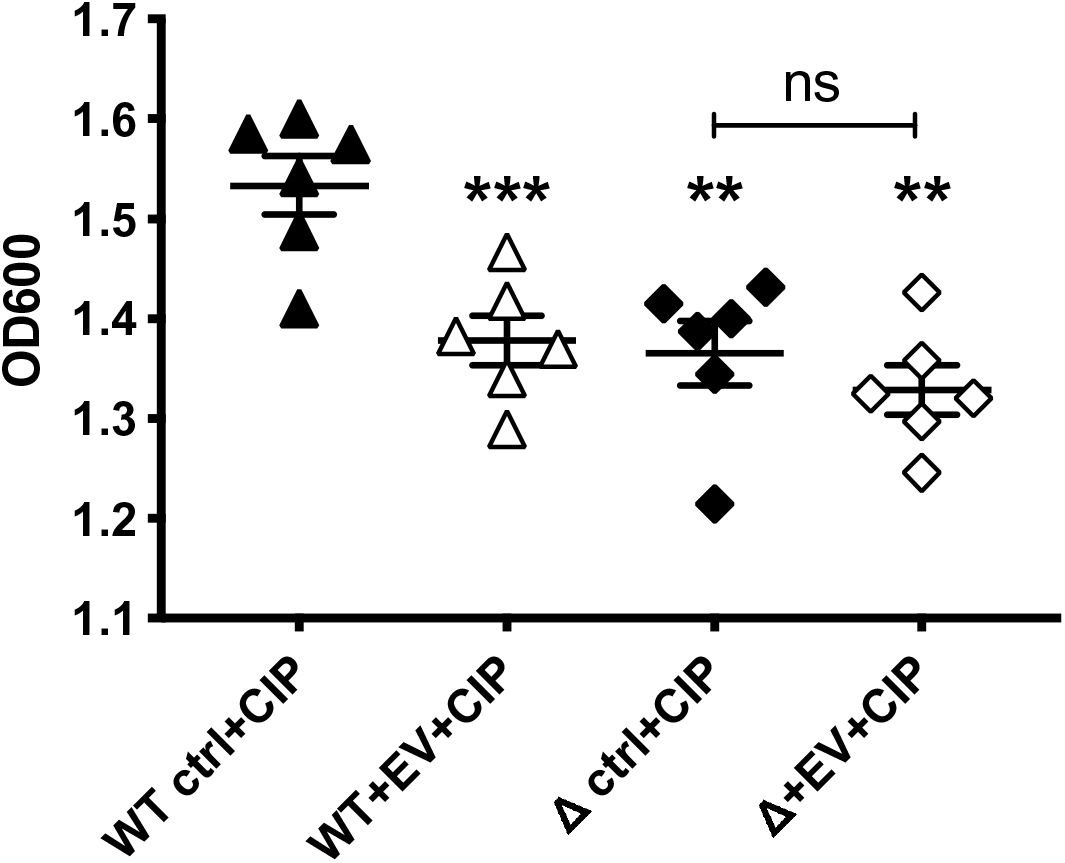
EVs increase *P. aeruginosa* ciprofloxacin sensitivity by targeting the fluoroquinolone efflux pump MexHI-OpmD. EVs decreased planktonic growth of wild type (WT) PAO1 in the presence of ciprofloxacin (0.03 µg/ml). EVs in the presence of ciprofloxacin (0.03 µg/ml) had no significant effect on the *mexHI-opmD* knockout strain (Δ+EV). Statistical significance was determined with repeated measures one-way ANOVA followed by Dunnett’s multiple comparisons test. EVs were isolated from six AEC donors and each data point is the mean of three technical replicates from each donor. ***P < 0.001 compared to WT ctrl, **P < 0.01 compared to WT ctrl, ns = not significant.

### Characterization of sRNA content of EVs

We previously found that EVs secreted by AEC deliver the miRNA let-7b-5p to *P. aeruginosa*, where it reduced biofilm formation and increased sensitivity to beta-lactam antibiotics by targeting key genes involved in biofilm formation and beta-lactam resistance (19). However, let-7b-5p alone did not reduce protein levels of MexHI-OpmD. To determine whether a different sRNA was responsible for the EV-mediated reduction in MexHI-OpmD protein levels and the resulting increased sensitivity to fluoroquinolone antibiotics, we re-analyzed our existing RNA-seq data (GSE174690) to identify sRNAs in EVs that are predicted to target MexHI-OpmD. The alignment-based characterization of EV sRNA content showed that tRNA and tRNA like fragments were the most abundant sequences in EVs, accounting for 65% of all aligned reads (Table 2). To gain a better understanding of the sRNA content of EVs at the sequence level, we generated count tables for unique sequences and filtered them for the most abundant unique sequences with at least 20 nucleotide length and at least 100 counts in each sample. The resulting 1346 sequences were aligned to homo sapiens using BLAST+ to obtain annotations for the 909 most abundant unique sequences with 100% identity and coverage (S2 Table). Collectively, these 909 unique sequences accounted for 60% of total reads detected in all 3 replicate samples. The sequences with the highest total count (listed at the top of S2 Table) include many tRNA and lncRNA fragments.

**Table 2.** Alignment-based characterization of EV sRNA content.

| RNA type | % of total reads |
| --- | --- |
| tRNA | 46.0 |
| tRNA like | 19.0 |
| RefSeq anti | 16.5 |
| RefSeq | 9.9 |
| rfam | 3.0 |
| rRNA | 1.8 |
| other ncRNA | 1.4 |
| piRNA | 1.1 |
| lncRNA | 0.7 |
| miRNA | 0.3 |
| lncRNA anti | 0.3 |
| other ncRNA anti | 0.1 |
| CDBox | 0.003 |

RNA types (column 1) and their relative abundance as a percentage of total reads (column 2).
EVs were derived from three individual AEC donors. tRNA = transfer RNA, RefSeq =
messenger RNA, anti = antisense, rfam = non-coding RNA annotated in Rfam database, rRNA
= ribosomal RNA, ncRNA = non-coding RNA, piRNA = Piwi-interacting RNA, lncRNA = long
non-coding RNA, miRNA = microRNA, CDBox = Small nucleolar RNA (snoRNA).

### Transfer of EV sRNAs to *P. aeruginosa*

We previously described the transfer of miRNAs secreted in AEC EVs to *P. aeruginosa* (19). To assess the transfer of other sRNAs besides miRNAs to *P. aeruginosa*, we re-analyzed RNA-seq data of EV-exposed *P. aeruginosa* and unexposed controls (GSE174710). We generated count tables of unique sequences from trimmed reads for individual samples - three unexposed controls and three samples of *P. aeruginosa* exposed to EVs secreted by different AEC donors. Reads were filtered to include only sequences with a length of at least 20 nucleotides that were detected in all three EV-exposed *P. aeruginosa* samples and none of the unexposed control samples. Filtering on these criteria yielded 1022 sequences, 15 of which were perfect matches to human sequences in a BLAST search. The length requirement of at least 20 nucleotides was chosen because shorter sequences tend to be less specific and map to many different regions in the genome. The 15 human EV sRNA sequences listed in Table 3 were detected in EV-exposed *P. aeruginosa*, but not in any of the unexposed control samples, suggesting that they are delivered to *P. aeruginosa* by EVs. Subsequently, we used IntaRNA to predict target genes of these 15 human EV sRNAs in *P. aeruginosa*.

**Table 3.**
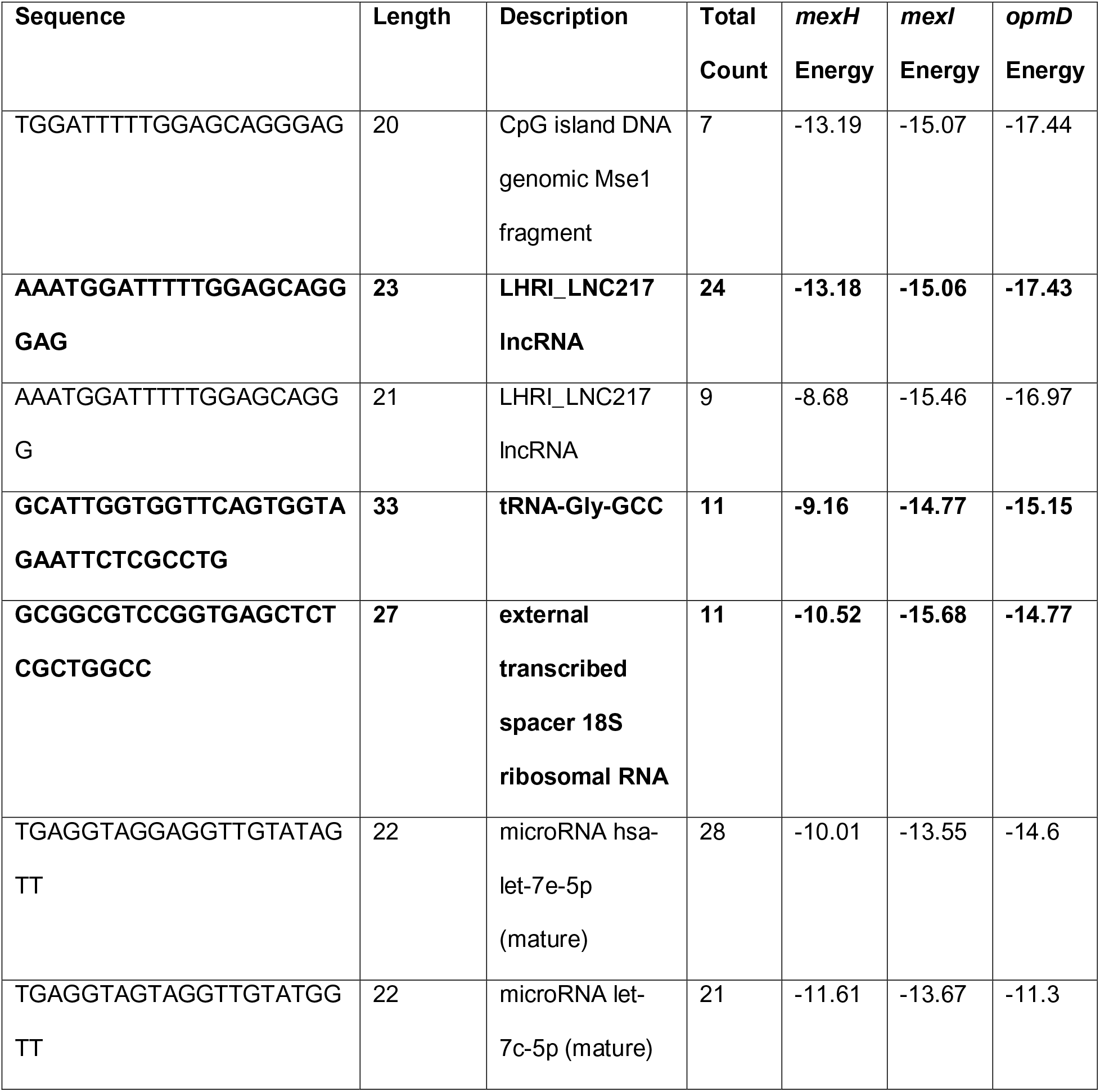
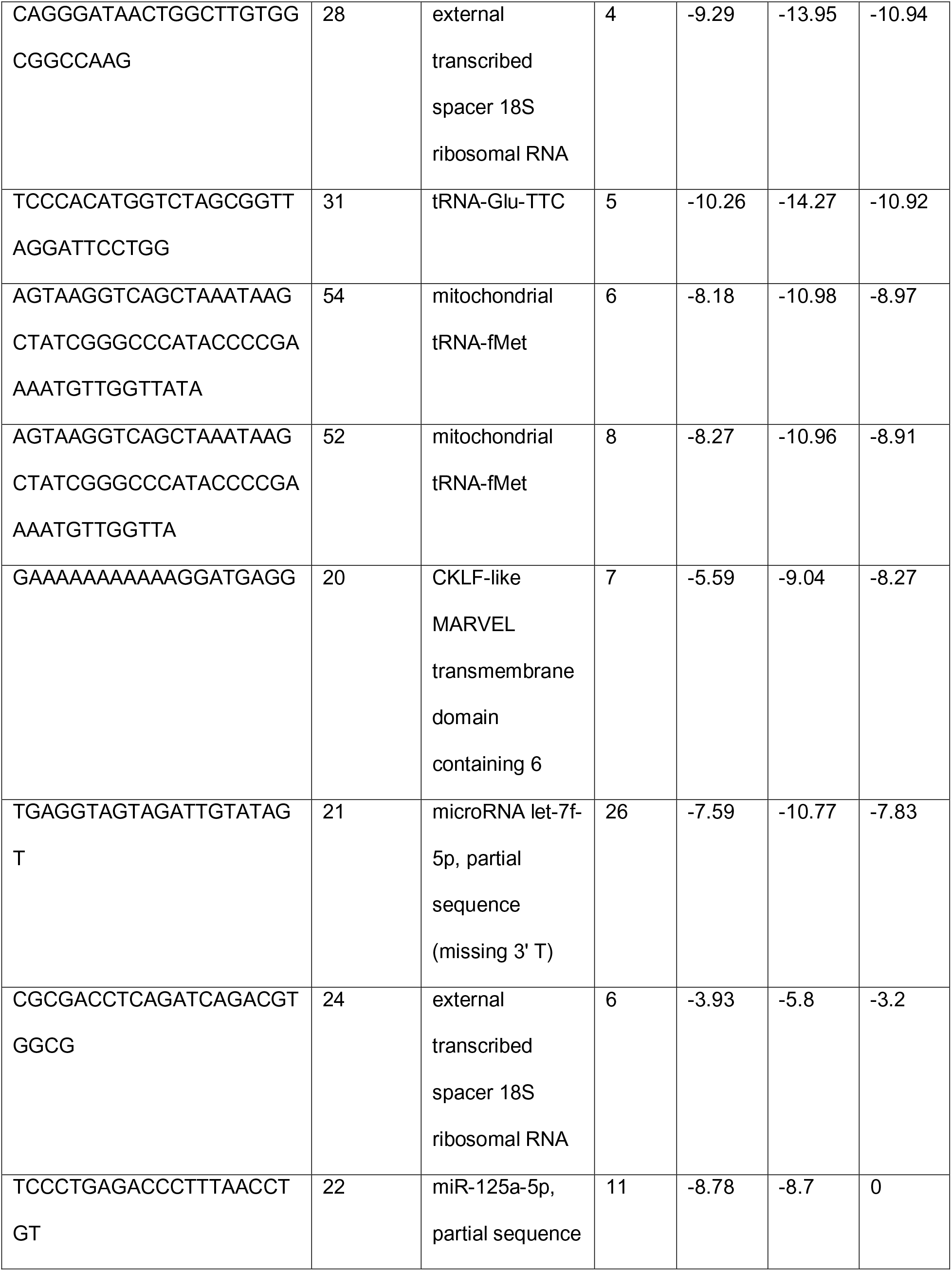
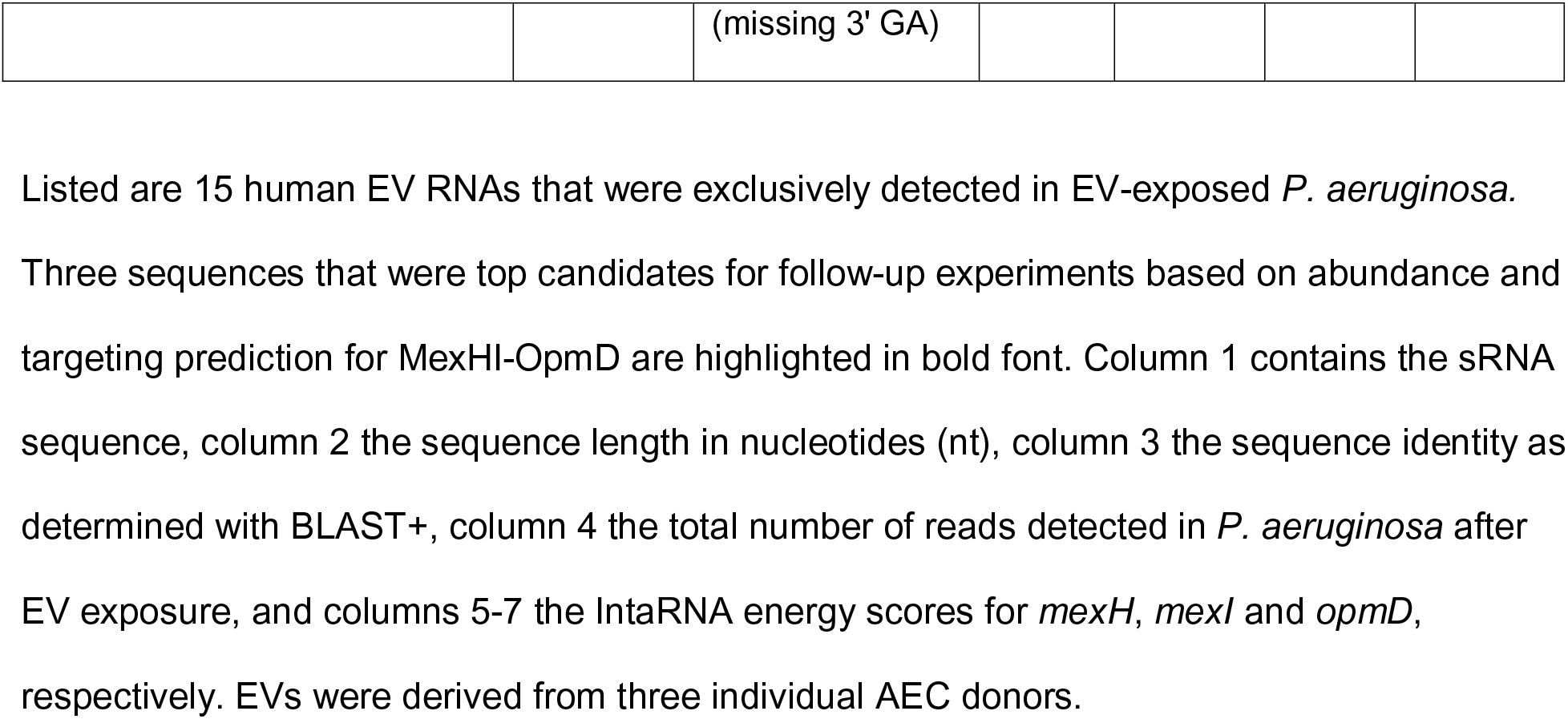
Human EV RNA fragments delivered to *P. aeruginosa*.

### Human EV sRNAs are predicted to target *P. aeruginosa* efflux pump *mexHI-opmD*

To predict which *P. aeruginosa* genes the 15 human sRNAs that were delivered to *P. aeruginosa* by EVs may target, the RNA-RNA interaction tool IntaRNA (50,51) was used to calculate binding energy scores. The more negative the energy score, the higher the likelihood of an RNA-RNA interaction. Table 3 lists the IntaRNA energy scores for each of the 15 EV sRNAs for predicted targeting of the fluoroquinolone efflux pump subunits *mexH, mexI*, and *opmD*. Three sRNAs with a high count in PA14+EV samples and the best (i.e., lowest) energy scores for targeting of *mexH, mexI*, and *opmD* were selected for follow-up validation experiments: a 23-nt lncRNA fragment, a 33-nt tRNA-Gly fragment and a 27-nt fragment of an 18S rRNA external transcribed spacer (highlighted in bold font in Table 3). As described next, of the three top candidate sRNAs based on bioinformatic predictions, the rRNA fragment was validated in wet lab experiments to increase the fluoroquinolone sensitivity of *P. aeruginosa* and to decrease MexHI-OpmD protein levels.

### A human rRNA fragment mediates the increased fluoroquinolone sensitivity of *P. aeruginosa* by targeting MexHI-OpmD

To test whether the 18S rRNA spacer fragment in EVs mediates the increased ciprofloxacin sensitivity of *P. aeruginosa* in response to EVs, we expressed the rRNA fragment (sequence listed in Table 3) in *P. aeruginosa* using a plasmid with an arabinose-inducible promoter and measured planktonic growth in the presence and absence of ciprofloxacin using the empty vector as the control strain (ctrl). We observed that the rRNA fragment significantly decreased planktonic growth of *P. aeruginosa* in the presence of ciprofloxacin (0.03 µg/ml, Fig 3A), while it did not significantly alter planktonic growth in the absence of ciprofloxacin (Fig 3B). Full 20-hour growth curves for the data in Fig 3A and 3B are provided in Fig S3. The rRNA fragment also significantly decreased the MIC of *P. aeruginosa* for ciprofloxacin (Fig 3C), and significantly reduced the number of live bacteria in the presence of ciprofloxacin (Fig 3D). In the presence of the rRNA fragment planktonic growth of *P. aeruginosa* was reduced by 76% compared to the empty vector control, the MIC of *P. aeruginosa* for ciprofloxacin was decreased two-fold and the number of live bacteria was reduced by 73%.

**Fig 3.**
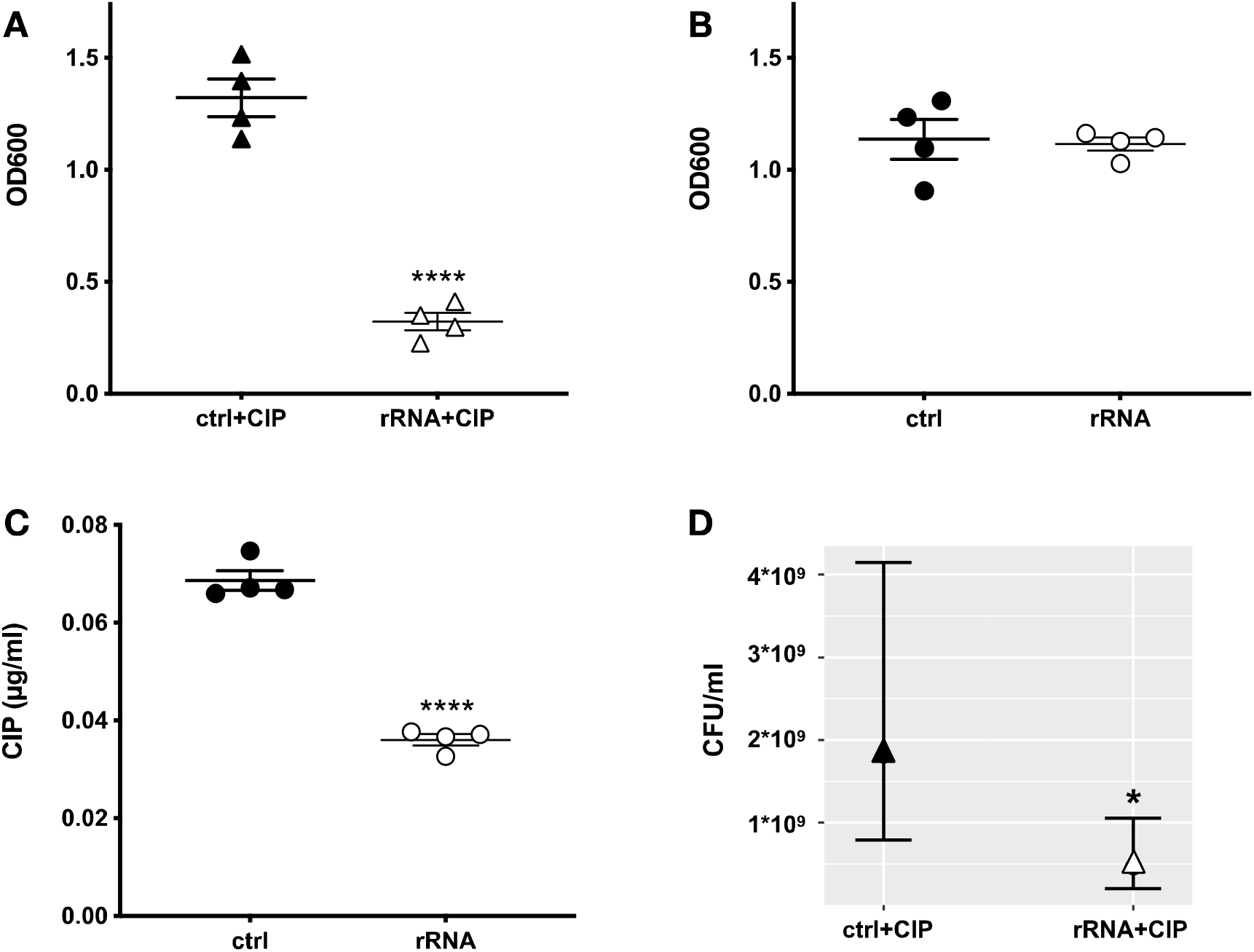
A human rRNA fragment increases the fluoroquinolone sensitivity of *P. aeruginosa*. Compared to the empty vector control (ctrl), the rRNA fragment significantly reduced planktonic growth (OD600) of PA14 in the presence of 0.03 µg/ml ciprofloxacin (CIP) and 100 mM arabinose after a 20-hour incubation (A) but did not significantly affect planktonic growth in the absence of ciprofloxacin (B). The rRNA fragment also significantly reduced the minimal inhibitory concentration (MIC) of ciprofloxacin (C) as well as the number of live bacteria in the presence of 0.03 µg/ml ciprofloxacin (D). Statistical significance was determined using unpaired t-tests for panels A-C with N = four independent experiments using different clones of each strain and each data point is the mean of three technical replicates and a negative binomial regression model for panel D with N = three independent experiments using different clones of each strain. Panel D shows the batch-corrected estimates from the regression model. CFU = colony forming units. ****P < 0.0001 compared to ctrl, *P < 0.05 compared to ctrl.

To validate that the rRNA fragment increased the ciprofloxacin sensitivity of *P. aeruginosa* by targeting the fluoroquinolone efflux pump MexHI-OpmD, we measured protein levels of MexHI-OpmD in the presence and absence of the rRNA fragment and found that compared to the empty vector control, the rRNA fragment decreased MexH by 53%, MexI by 26% and OpmD by 52% (Table 4). The decrease in the protein levels of MexH and OpmD was statistically significant, and there was a trend for the rRNA fragment to decrease MexI protein levels that did not quite reach statistical significance. Normalized peak intensities for each sample set as well as average log2 fold changes and p-values for all 1911 detected proteins in *P. aeruginosa* are provided as S3 Table. Taken together, these experiments demonstrate that the rRNA fragment increased the ciprofloxacin sensitivity of *P. aeruginosa* by targeting the fluoroquinolone efflux pump MexHI-OpmD.

**Table 4.** A human rRNA fragment significantly reduces protein levels of MexH and OpmD.

| Uniprot ID | Locus | Name | Product | log2 FC | % decrease | p-value |
| --- | --- | --- | --- | --- | --- | --- |
| A0A0H2ZGA2 | PA14_09530 | MexH | RND efflux membrane fusion protein | -1.09 | -53 | 0.0003 |
| A0A0H2ZGB3 | PA14_09520 | MexI | RND efflux transporter | -0.43 | -26 | 0.1344 |
| A0A0H2ZF64 | PA14_09500 | OpmD | outer membrane protein | -1.07 | -52 | 0.0107 |

Protein level changes in a *P. aeruginosa* strain induced with 100 mM arabinose to express the rRNA fragment compared to the empty vector control strain. FC = fold change, N = three different clones of each strain. Statistical significance was determined with QPROT.

## Discussion

We report that AECs secrete EVs containing lncRNA, tRNA and 18S rRNA external transcribed spacer fragments that are transferred to *P. aeruginosa*, and that EVs increased the fluoroquinolone sensitivity of *P. aeruginosa* by reducing protein levels of the drug efflux pump MexHI-OpmD. An 18S rRNA external transcribed spacer fragment by itself significantly reduced the protein levels of MexH and OpmD, thereby increasing antibiotic sensitivity to ciprofloxacin. The finding that AEC EVs contain rRNA fragments is consistent with prior characterizations of the sRNA content of EVs secreted by airway epithelial cells (40,41). It has been previously described that rRNA fragments are produced in a controlled manner that is population-, sex- and tissue-specific, suggesting they constitute another class of small non-coding regulatory RNAs (52,53). Several reports predict and speculate about a regulatory function of rRNA derived fragments (52,54,55), with some studies providing evidence for the regulation of various biological functions by rRNA fragments. Chen et al. report RNAi knockdown of an endogenous 20-nt 28S rRNA fragment in the human non-small cell lung carcinoma cell line H1299 induced apoptosis and inhibited cell proliferation (56), while Wei et al. found that endogenous rRNA fragments detected in human and murine liver regulate glucose metabolism in the murine hepatoma cell line Hepa 1-6 and are differentially expressed in the livers of diabetic mice (57). Moreover, several studies show co-immunoprecipitation of eukaryotic rRNA fragments with Argonaute proteins (54,57,58), further corroborating our conclusion that rRNA fragments are involved in the regulation of gene expression. Most of the previously described rRNA fragments are derived from 5.8S, 18S or 28S rRNA as well as the internal transcribed spacer and, in fact, some sequences that had been previously classified as miRNAs or piRNAs are in fact rRNA-derived (53). There is very limited knowledge about the existence and function of fragments derived from the rRNA external transcribed spacer. Lee et al. describe upregulation of rRNA external transcribed spacer fragments in *Neurospora crassa* in response to DNA damage (59), while Gupta et al. found that fragments derived from an rRNA external transcribed spacer are upregulated in the parasitic protist *Entamoeba histolytica* in response to growth stress, suggesting that these fragments play a regulatory role in the stress response (60). However, direct evidence for the regulation of gene or protein expression by rRNA external transcribed spacer fragments has not been published. In addition, to our knowledge there are no studies showing cross kingdom transfer of rRNA fragments from human EVs to a bacterial pathogen, nor is anything known about the effect of human rRNA fragments on bacterial gene expression. Thus, to our knowledge, our data represent the first report of an rRNA fragment secreted by a eukaryotic cell that regulates protein expression and alters the phenotype of a prokaryotic organism. The exact mechanism of action and potential protein binding partners of rRNA fragments in prokaryotes are beyond the scope of this work and need to be elucidated in future studies.

We found that some, but not all, of the most abundant sequences in EVs can be detected in *P. aeruginosa* following EV exposure. However, some human EV sRNA sequences detected in EV-exposed *P. aeruginosa* are not among the most abundant EV sequences, suggesting differential stability in *P. aeruginosa*. While RNA contained within EVs is protected from degradation by extracellular RNases, once the EV RNA cargo is delivered to *P. aeruginosa*, RNA that is not protected by binding to protein or *P. aeruginosa* mRNA targets may be more prone to degradation, explaining the discrepancy between abundance in EVs compared to the abundance after delivery to *P. aeruginosa* for some of the sRNAs.

One limitation of our study is that we did not assess other AEC EV cargo such as DNA, protein, lipids, or metabolites, which could contribute to EV induced sensitivity to ciprofloxacin. However, our experiment expressing the rRNA fragment in *P. aeruginosa* demonstrates that the rRNA fragment alone is sufficient to reduce the abundance of MexH and OpmD and to increase the sensitivity to ciprofloxacin.

Another limitation is that due to redundancy of fluoroquinolone efflux pumps in *P. aeruginosa*, the suppression of MexHI-OpmD only leads to a partial effect on fluoroquinolone sensitivity. MexEF-OprN serves as another fluoroquinolone efflux pump in *P. aeruginosa* (61), and there are several additional efflux pumps that can serve this function (62), which helps explain why deletion of MexHI-OpmD does not result in a more drastic increase in fluoroquinolone sensitivity (Fig 2). On the other hand, as shown in Fig 2, the downregulation of MexHI-OpmD by EVs is enough to reduce fluoroquinolone sensitivity to a level achieved with a complete deletion of the efflux pump. Moreover, we found that expressing the rRNA fragment by itself led to a larger reduction in *P. aeruginosa* planktonic growth in the presence of ciprofloxacin than EVs, and that while both EVs and the rRNA fragment alone led to a protein level reduction of the fluoroquinolone efflux pump MexHI-Opm, the rRNA fragment significantly suppressed additional proteins involved in mediating fluoroquinolone resistance, including MexA, GyrA, and ParC (62,63). This may explain the increased ability of the rRNA fragment to reduce planktonic growth in the presence of ciprofloxacin compared to EVs, which contain additional cargo that may counteract the effect of the rRNA fragment.

*P. aeruginosa* in the lung is exposed to EVs secreted by other cell types in addition to those secreted by AEC, most notably immune cells like neutrophils and macrophages, whose EV RNA content may include the rRNA fragment in our study as well as other sRNAs that may affect antibiotic resistance of *Pseudomonas*. Therefore, a final limitation is that our study cannot elucidate the cumulative effect that EV sRNAs from multiple cell types would have on *P. aeruginosa* in the lungs. However, re-analysis of publicly available data (GSE126051) from a study characterizing sRNAs in human plasma (64) revealed that rRNA fragments that were identical in length and sequence to the rRNA fragment described in this study were present in plasma of healthy control subjects. Moreover, identical rRNA fragments were detected in EVs isolated from human bronchoalveolar lavage fluid (BALF; Dr. Alix Ashare, personal communication). These observations reveal the biological significance of our study since our rRNA fragment was found in clinical samples and is not limited to our model system of primary AECs. So far, there are no published studies that comprehensively characterize the sRNA content of EVs isolated from human BALF, and, as noted in a recent review by Zareba et al., future in-depth RNA-seq profiling of distinct EV populations in BALF will elucidate their role in disease pathogenesis and provide insight into treatment strategies (65). The important questions of whether *P. aeruginosa* exposure alters the sRNA content of AEC EVs and whether there is a difference in the sRNA cargo of EVs secreted by WT and CF AEC need to be addressed in future studies.

To assess whether EV sRNA targeting of efflux pumps is limited to *P. aeruginosa* or extends to efflux pump orthologs in other common lung pathogens we utilized IntaRNA to predict targeting of *Burkholderia cenocepacia* (which has orthologs for *mexH, mexI* and *opmD*) and *Staphylococcus aureus* (which has an ortholog for *mexI*). Our candidate rRNA spacer fragment is predicted to target orthologs of *mexI* and *opmD* in *Burkholderia cenocepacia*, suggesting that AEC EV sRNAs are likely to affect other lung pathogens beyond *P. aeruginosa* and warrant future analysis beyond the scope of the current study.

In summary, our study provides the first evidence for transfer of lncRNA, tRNA and rRNA fragments from eukaryotic EVs to a prokaryote and provides direct evidence for regulation of the prokaryote by a eukaryotic rRNA fragment, resulting in subsequent phenotypic alterations such as increased fluoroquinolone sensitivity. The development of new treatment approaches utilizing a combination of human rRNA fragments and antibiotics in nanoparticles or EVs may benefit individuals with chronic antibiotic-resistant *P. aeruginosa* infections.

## Materials and Methods

### Culture of airway epithelial cells and EV isolation

De-identified primary human AEC were obtained from Dr. Scott Randell (University of North Carolina, Chapel Hill, NC) and cultured in BronchiaLife basal medium (Lifeline Cell Technology, Frederick, MD, Cat. # LM-0007) supplemented with the BronchiaLife B/T LifeFactors Kit (Lifeline Cell Technology, Cat. # LS-1047) as well as 10,000 U/ml Penicillin and 10,000 μg/ml Streptomycin (Sigma-Aldrich, Cat. # P4333), as previously described (66). EVs were isolated from AEC culture supernatants (passages 4-8) with the ExoQuick-TC EV isolation kit (System Biosciences, Palo Alto, CA, Cat. # EXOTC50A-1) and characterized as previously described (67).

### Bacterial strains and culture

For all experiments, *P. aeruginosa* strains PA14, PAO1, and the mucoid CF clinical isolate SMC1585 (49) were cultured in Luria broth (LB, Thermo Fisher Scientific, Waltham, MA). A deletion mutant for the fluoroquinolone efflux pump *mexGHI-*opmD in the PAO1 background (strain DW101, PAO1ΔGHID), as well as the matching parental PAO1 wild type strain, were generously provided by Dr. Helen Zgurskaya from the University of Oklahoma (61). Deletion of *opmD* in the PAO1*ΔmexGHI-*opmD strain was confirmed by PCR. Briefly, RNA was isolated from PAO1*ΔmexGHI-opmD* and the parental PAO1 wild type strain using the miRNeasy Mini Kit (Qiagen, Germantown, MD, Cat. # 217004), cDNA was synthesized with the Invitrogen RETROscript Reverse Transcription Kit (Thermo Fisher Scientific Cat. # 10585595), and the presence of *opmD* was interrogated by PCR amplification using the following primers: forward 5’-CCTGGTGGAGTTTCTTCGAC-3’ and reverse 5’-CGTCGTAGTCCAGCTGTTGT-3’. *P. aeruginosa* strains expressing either the 18S rRNA spacer fragment under the control of an arabinose-inducible promoter or the empty vector control were produced by transforming PA14 with a pMQ70 expression vector (68) containing either the rRNA fragment sequence (listed in Table 3) as an insert, or no insert (control). Strains were cultured with 300 µg/ml carbenicillin (Sigma-Aldrich, Cat. # C1389) to select for bacteria containing the plasmid and 100 mM L-(+)-arabinose (Sigma-Aldrich, St. Louis, MO, Cat. # A3256) was added during experiments to induce expression of the rRNA fragment.

### Bacterial growth curves and CFU count

The minimal inhibitory concentration (MIC) of the fluoroquinolone antibiotic ciprofloxacin was determined by incubation of each *P. aeruginosa* strain with 8 different concentrations of ciprofloxacin (0-0.25 µg/ml) in triplicate for 20 hours in a plate reader at 37°C. The OD600 was recorded every 15 minutes and the MIC for ciprofloxacin was calculated by fitting the growth curve data with a Gompertz function, as previously described (69,70). At the beginning of the 20-hour growth curves, 5000 colony forming units (CFUs) of *P. aeruginosa* were exposed to 5×10^9^ EV/ml, corresponding to a concentration of EVs measured in AEC culture supernatants by us, as well as in human bronchoalveolar lavage fluid (42,71). During exposure of *P. aeruginosa* to EVs, ciprofloxacin (Sigma-Aldrich, Cat. # 17850-5G-F) was present in the culture medium at a concentration equivalent to one-third to one-half the MIC of a given strain, as indicated in the figure legends. CFU/ml for viable planktonic bacteria were obtained by dilution plating of supernatants.

### Proteomics of *P. aeruginosa*

1.5*10^7^ CFUs of PA14 were exposed to 1.5*10^10^ EV isolated from AEC from three donors or vehicle controls for a PA14:EV ratio of 1:1000 and incubated for 15 h at 37°C at 225 rpm. Samples were processed and TMT labeled as described previously (19). All six samples were run in duplicate, peak intensities from duplicate runs were averaged and filtered to retain 1619 proteins that were detected in all replicate samples. Differentially abundant proteins were identified using QPROT (72). The mass spectrometry proteomics data have been deposited to the ProteomeXchange Consortium (73) via the MassIVE (74) partner repository with the dataset identifier PXD033213.

In addition, following overnight cultures with 300 µg/ml carbenicillin to select for bacteria with the plasmid containing the rRNA fragment or the empty plasmid, 3 different clones of the PA14 strains expressing either the rRNA fragment or the empty vector control were cultured with 100 mM arabinose for 15 h at 37°C at 225 rpm to induce expression of the rRNA fragment. Cell pellets were washed with PBS, lysed and proteins (200 µg of each sample) were purified by acetone/trichloroacetic acid precipitation overnight at -20°C. After washing the pellet with ice-cold acetone, the resulting protein pellet was resuspended in 50 μL of 8 M urea in 400 mM ammonium bicarbonate, pH 7.8, reduced with 4 mM dithiothreitol at 50°C for 30 minutes, and cysteines were alkylated with 18 mM iodoacetamide for 30 minutes. The solution was then diluted to <2 M urea and trypsin (Promega, Madison, WI) was added at a final trypsin/protein ratio of 1:100 prior to overnight incubation at 37°C. The resulting peptides were desalted using solid-phase extraction on a C18 Spin column and eluted in 240 μL of 80% acetonitrile in 0.1% formic acid. All samples were analyzed in technical duplicates by LC-MS/MS using a nanoElute coupled to a timsTOF Pro Mass Spectrometer (Bruker Daltonics). Two hundred nanogram of each digested peptide sample was loaded on a capillary C18 column (25 cm length, 75 μm inner diameter, 1.6 μm particle size, 120 Å pore size; IonOpticks). The flow rate was kept at 300 nL/min. Solvent A was 0.1% FA in water, and Solvent B was 0.1% FA in ACN. The peptides were separated on a 100 min analytical gradient from 2% ACN/0.1% FA to 35% ACN/0.1% FA for a total of 120 min gradient. The timsTOF Pro was operated in the PASEF mode. MS and MS/MS spectra were acquired from 100-1700 m/z. The inverse reduced ion mobility 1/K_0_ was set to 0.60−1.60 V.s/cm^2^ over a ramp time of 100 ms. Data-dependent acquisition was performed using 10 PASEF MS/MS scans per cycle with a near 100% duty cycle. The resulting protein tandem MS data was queried for protein identification and label-free quantification against the Uniprot *Pseudomonas aeruginosa* strain UCBPP-PA14 database (proteome ID UP000000653, retrieved on Feb 2, 2022) using MaxQuant v2.0.3.1. The following modifications were set as search parameters: peptide mass tolerance at 20 ppm, trypsin digestion cleavage after K or R (except when followed by P), 2 allowed missed cleavage sites, carbamidomethylated cysteine (static modification), and oxidized methionine, deaminated asparagine/glutamine, and protein N-term acetylation (variable modification). Search results were validated with peptide and protein FDR both at 0.01. LFQ intensities were averaged across technical replicates, filtered to retain 1911 proteins detected in at least two replicate samples, and normalized prior to QPROT analysis (72). The mass spectrometry proteomics data have been deposited to the ProteomeXchange Consortium (73) via the PRIDE (75) partner repository with the dataset identifier PXD033165.

### Characterization of EV sRNA content

Previously, we characterized the miRNA content of AEC EVs (19); here, we focus on the EV sRNA content beyond miRNAs. The raw data as well as count tables of aligned reads provided by System Biosciences can be accessed through NCBI’s Gene Expression Omnibus (GEO Series accession number GSE174690). In addition to the alignment-based characterization we also performed a sequence-based characterization of EV sRNA content. Adaptor trimming of raw reads was performed with fastp version 0.20.0 (76). Trimmed reads from EVs secreted by three donors of AEC yielded close to 500,000 unique sequences, about 40,000 of which were detected in all three biological replicates. Count tables with the number of reads per sample for each unique sequence were generated using the Unix command “cat fastp.fastq.gz | gunzip | awk ‘(NR%4==2)’ | sort | uniq -c | sort -k 1 -n”. To determine the origin of the most abundant unique EV sRNA sequences, we ran command line BLAST+ (77) for 1346 sequences that had a minimum of 100 counts per sample and a length of at least 20 nucleotides. Arguments to the blastn function included -taxids 9606 (homo sapiens), -perc_identity 100 (requiring a perfect match), -qcov_hsp_perc 100 (requiring 100% query coverage) and -task blastn-short (to adjust for a short input sequence). Genbank accessions returned by BLAST were annotated using the R package rentrez (78). Subsequently, RNA-seq samples from EV-exposed *P. aeruginosa* were used to assess whether human sRNAs are transferred to *P. aeruginosa* by EVs.

### RNA-seq to detect transfer of EV RNA to *P. aeruginosa*

Existing RNA-seq data of *P. aeruginosa* exposed to PBS vehicle control in triplicate or to EVs from 3 AEC donors (19) was re-analyzed to identify EV-mediated transfer of sRNA other than miRNAs to *P. aeruginosa*. The raw data can be accessed through NCBI’s Gene Expression Omnibus (GEO Series accession number GSE174710). For each sample, trimmed reads were compiled into count tables of unique sequences and samples were filtered for unique sequences of at least 20 nucleotides in length that were contained in all three EV-exposed *P. aeruginosa* samples, but none of the control samples. The origin of 1022 unique sequences matching these criteria was determined using BLAST+ (77), as described above, and included a separate alignment to -taxids 287 (*P. aeruginosa*). Genbank accessions returned by BLAST were annotated using the R package rentrez (78). For EV RNA sequences that were of human origin, we generated IntaRNA targeting predictions (50,51) for *P. aeruginosa* genes, as described below.

### IntaRNA target predictions

We used the IntaRNA algorithm (50,51) to predict whether human sRNAs transferred to *P. aeruginosa* by EVs might regulate *P. aeruginosa* gene expression by targeting bacterial mRNAs. *P. aeruginosa* UCBPP-PA14 reference sequences and standard parameters were used to obtain energy scores as a measure of the likelihood of RNA-RNA interaction based on sequence similarity. The more negative the energy score, the higher the likelihood of interaction. Three top candidate RNA fragments derived from a lncRNA, a tRNA, and an rRNA were selected based on predicted targeting of MexHI-OpmD and abundance in *P. aeruginosa* following EV exposure. To assess whether targeting predictions extend beyond *P. aeruginosa* to other common lung pathogens, we ran IntaRNA to predict targeting of MexHI-OpmD orthologs by our top candidate RNA fragments in *Burkholderia cenocepacia* J2315, a multidrug-resistant CF clinical isolate, and *Staphylococcus aureus* COL, a MRSA clinical isolate.

### Statistical Analysis

Data were analyzed with Prism 8 for macOS (version 9.3.1, Graphpad, San Diego, CA) and the R software environment for statistical computing and graphics version 4.1.1 (79) using appropriate statistical methods, as indicated in the figure legends. Statistically significant differences between bacterial growth curves were determined with permutation tests using the compareGrowthCurves function from the R package statmod (version 1.4.36) (80,81).

## Acknowledgements

The authors would like to thank Dr. Helen Zgurskaya (University of Oklahoma) for providing the PAO1 deletion mutant for the fluoroquinolone efflux pump *mexGHI-*opmD as well as the matching parental PAO1 wild type strain. Mass Spectrometry analyses of the rRNA fragment expressing and empty vector strain of *P. aeruginosa* were performed by the Mass Spectrometry Technology Access Center at McDonnell Genome Institute (MTAC@MGI) at Washington University School of Medicine. This work was supported by funding from the Cystic Fibrosis Foundation (https://www.cff.org/) to B.A.S. (STANTO19G0, STANTO20P0, and STANTO19R0) and GREENE21G0 to D.A.H. and from the National Institutes of Health (https://www.nih.gov/) to B.A.S (P30-DK117469 and R01HL151385) and to S.A.G. (R01GM122846). The funders had no role in study design, data collection and analysis, decision to publish, or preparation of the manuscript.

## Supporting information

**S1 Fig.**
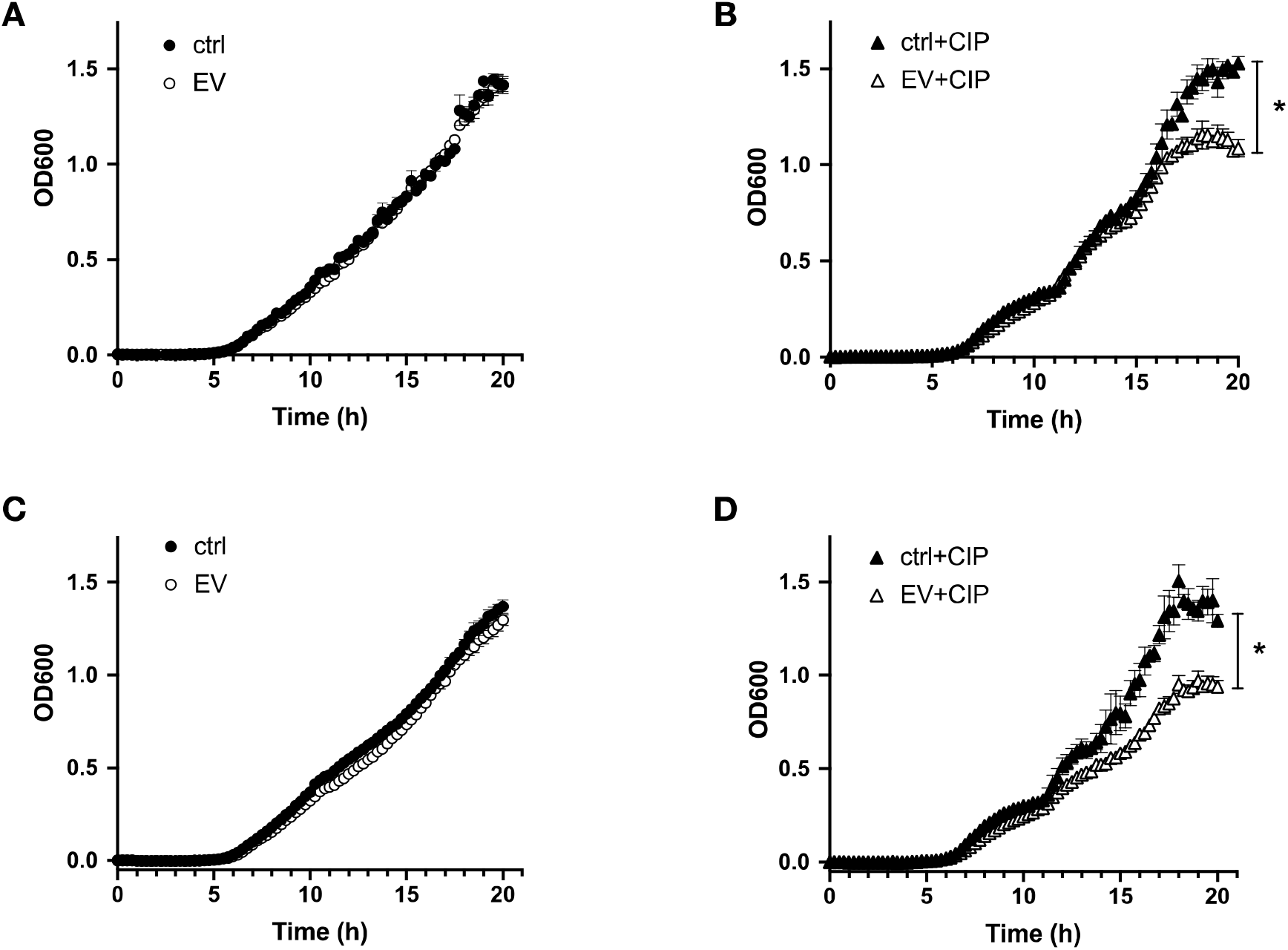
EVs increase the fluoroquinolone sensitivity of *P. aeruginosa*. (A) 20-hour planktonic growth curves (OD600) of PA14 exposed to EVs or vehicle control (ctrl) in the absence of ciprofloxacin. There was no significant difference between the growth curves for the last 4 hours of growth. (B) 20-hour planktonic growth curves (OD600) of PA14 exposed to 0.02 µg/ml ciprofloxacin and EVs or 0.02 µg/ml ciprofloxacin and vehicle control (ctrl). There was a significant difference between the growth curves for the last 4 hours of growth. (C) 20-hour planktonic growth curves (OD600) of clinical isolate SMC1585 exposed to EVs or vehicle control (ctrl) in the absence of ciprofloxacin. There was no significant difference between the growth curves for the last 4 hours of growth. (D) 20-hour planktonic growth curves (OD600) of clinical isolate SMC1585 exposed to 0.02 µg/ml ciprofloxacin and EVs or 0.02 µg/ml ciprofloxacin and vehicle control (ctrl). There was a significant difference between the growth curves for the last 4 hours of growth. Each data point shows the mean +/- SEM of EVs isolated from four AEC donors (or matching controls), averaged across three technical replicates. Statistical significance was determined using permutation tests. * P < 0.05.

**S2 Fig.**
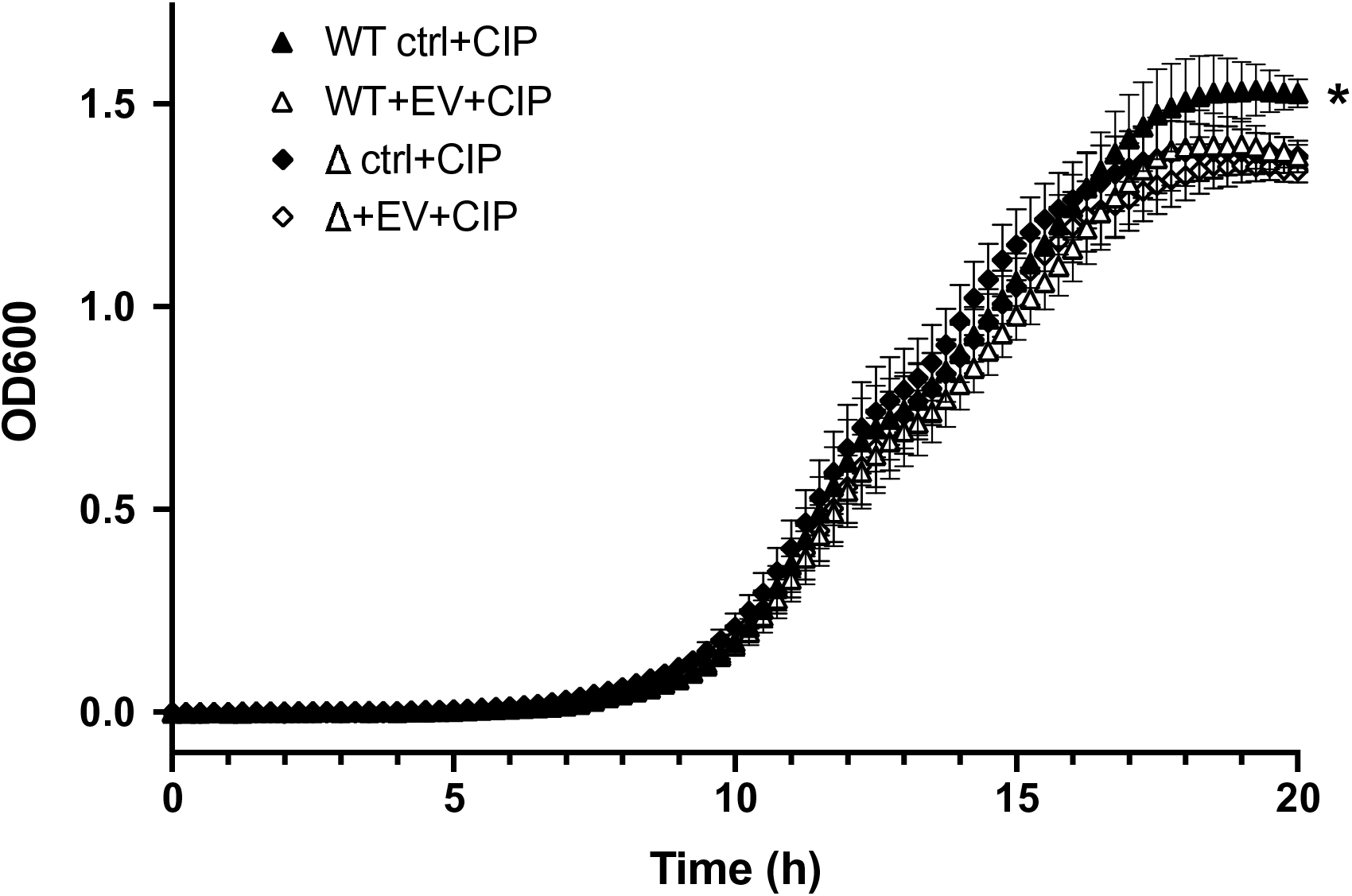
EVs increase *P. aeruginosa* ciprofloxacin sensitivity by targeting the fluoroquinolone efflux pump MexHI-OpmD. 20-hour planktonic growth curves (OD600) of PAO1 wild type (WT) and the PAO1 *mexHI-opmD* knockout strain (Δ) exposed to 0.03 µg/ml ciprofloxacin (CIP) in the presence and absence of EVs. There was a significant difference between WT ctrl+CIP and all other growth curves for the last hour of growth, while here was no significant difference between the other three growth curves. Each data point shows the mean +/-SEM of EVs isolated from six AEC donors (or matching controls), averaged across three technical replicates. Statistical significance was determined using permutation tests. * P < 0.05.

**S3 Fig.**
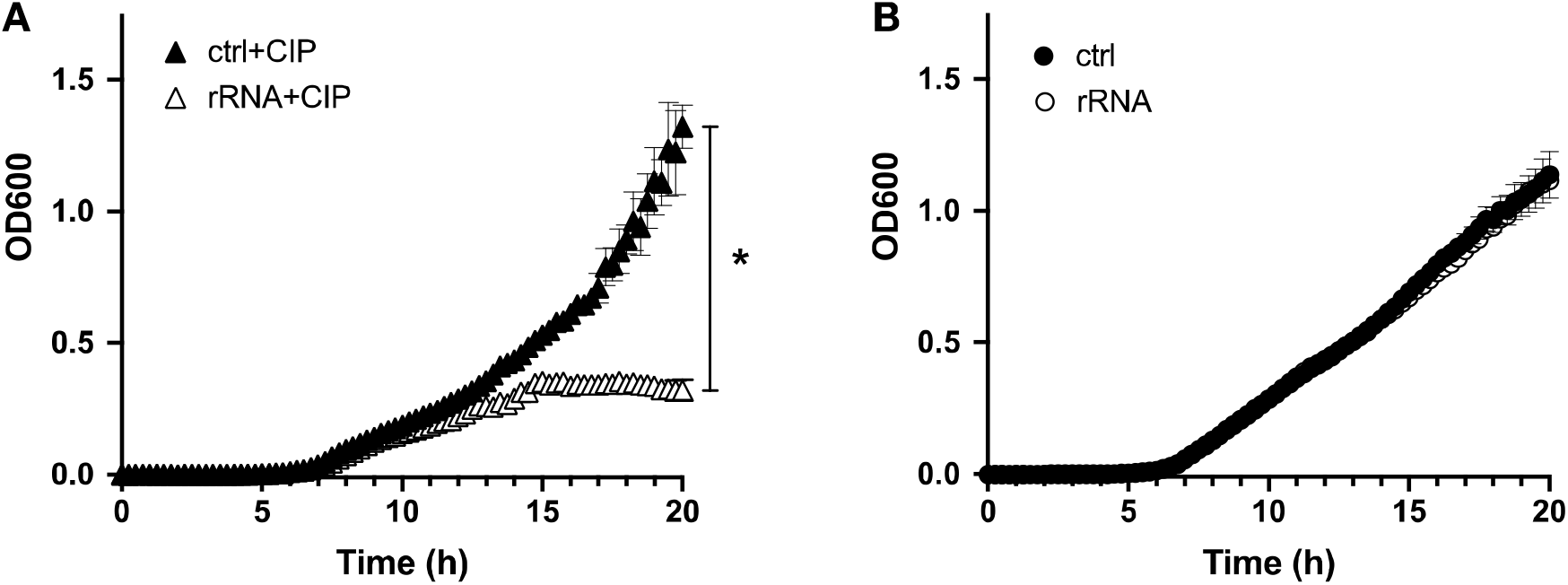
A human rRNA fragment increases the fluoroquinolone sensitivity of *P. aeruginosa*. (A) 20-hour planktonic growth curves (OD600) of PA14 expressing the rRNA fragment or empty vector control (ctrl) in the presence of 0.03 µg/ml ciprofloxacin (CIP) and 100 mM arabinose. There was a significant difference between the growth curves for the empty vector control (ctrl) and the rRNA expressing strain. (B) There was no significant difference in the 20-hour planktonic growth curves (OD600) of PA14 expressing the rRNA fragment or empty vector control in the absence of ciprofloxacin. Each data point shows the mean +/-SEM from four different clones of each strain averaged across three technical replicates. Statistical significance was determined using permutation tests. * P < 0.05.

**S1 Table. Proteomics data of EV-exposed *P. aeruginosa* and unexposed controls**. Normalized peak intensities for each sample as well as average log2 fold changes and p-values for 1619 proteins that were detected in duplicate runs of all six samples. EVs were derived from three individual AEC donors.

**S2 Table. Origin of most abundant sRNAs from the sequence-based characterization of EV sRNA content**. 909 most abundant unique sequences with at least 100 counts per sample accounted for 60% of total reads for sequences detected in all 3 replicate samples. Listed are the nucleotide sequence (column 1), sequence length (column 2), number of reads in EVs (column 3) and origin of sequence as determined with BLAST+ (column 4). EVs were derived from three individual AEC donors.

**S3 Table. Proteomics data of *P. aeruginosa* expressing the rRNA fragment compared to empty vector controls**. Normalized peak intensities for each sample as well as average log2 fold changes and p-values for 1911 proteins that were detected in at least two of three replicate samples.

